# Improved gene targeting in vivo using EoHR, a small molecule inhibitor of 53BP1

**DOI:** 10.1101/2024.05.30.596757

**Authors:** Sarah J. Spencer, Finn O’donoghue, Angelina Ristovski, Astrid Glaser, Subashani Maniam, Mitra Mohsenipour, Alita Soch, Andrew J. Deans, Stephen Headey

## Abstract

Precise genome editing by programmable nucleases such as Cas9 has revolutionised medical research by enabling the creation of gene-edited or knock-in mouse models of disease. However, a major limitation of the approach is the inefficient process of homology driven recombination (HDR) from an exogenous DNA repair template. This is because error-prone, 53BP1-dependent non-homologous end joining (NHEJ) predominates at Cas9-induced double strand breaks (DSBs). Here we report the validation of a cell-permeable and non-toxic inhibitor of 53BP1 called EoHR. In vitro, EoHR prevents 53BP1 binding to dimethylated lysine 20 on histone H4, a marker of DSBs. In cells, EoHR prevents localisation of 53BP1 at nuclease mediated DSBs and promotes HDR at a Cas9-induced break. When tested in vivo during mouse Cas9-mediated gene editing, inclusion of EoHR at the time of microinjection more than doubled the recovery of correctly edited mice and halved the time to project success. Our work shows that inhibition of 53BP by EoHR is a simple and robust way to increase HDR at Cas9 breaks that can also dramatically increase the success rates of animal model generation.

## Introduction

There are three principal uses of CRISPR in generating animal models: gene knock-out, gene knock-in, and gene editing, such as introducing or correcting a disease-causing point mutation. Knock-out CRISPR is relatively efficient as it exploits error-prone NHEJ, the dominant DSB repair pathway (Cong et al., 2013). In contrast, specific DNA editing with Cas9 or its variants requires either prime editing (Anzalone et al., 2019) to avoid creating DSBs or homology-driven recombination (HDR) CRISPR to mediate DSB repair. HDR CRISPR is generally the method of choice for generating complex alleles in mice. In the repair of spontaneous DSBs homologous recombination (HR) utilises the sister chromosome as the template for DNA repair and is thus error-free (Her and Bunting, 2018). When HDR CRISPR is used to introduce a gene, or specifically edit DNA, in addition to the single guide RNA (sgRNA) and Cas9, a DNA repair template must be provided that contains the desired genetic insert and has DNA arms homologous to the host DNA adjacent to the Cas9-induced DSB (Harms et al., 2014). This exogenous DNA substitutes for the sister chromosome to act as the repair template for HDR incorporation of the desired mutation. The difficulty for precise HDR CRISPR editing is that HR is tightly regulated and is usually restricted to late S and G2 phases of the cell cycle when the sister chromosomes are available (Her and Bunting, 2018).

Considerable variability in success is also apparent depending on the locus selected for insertion. Some loci, such as the Rosa locus (Friedrich and Soriano, 1991) are much more amenable to HDR CRISPR insertions, although efficiency is still low. However, to recapitulate a human model often requires editing or inserting a gene at a specific chromosomal location. Even with recent advances to our understanding in sgRNA design, the overall efficiency of HDR CRISPR is poor; when targeting biologically appropriate loci approximately 3% of mouse pups on average will have the correct DNA insertion, as evidenced in this study. Similar success rates of 0-10% are also reported for knock-in rabbits (Song et al., 2016). Thus, generating a single founder knock-in animal using HDR CRISPR is usually a slow and expensive process with a high failure rate.

Compounds tested against a variety of targets to enhance HDR CRISPR *in vitro* have shown mixed results depending on the cell line selected (Riesenberg and Maricic, 2018). Variable activity of the Fanconi anaemia pathway in single-strand template repair might, to some extent, explain the cell-type variability (Richardson et al., 2018). Crucially, no compound is reliably validated *in vivo*. Initial attempts at HDR CRISPR enhancement with the putative RAD51 stimulator, RS-1 (Jayathilaka et al., 2008) showed promise *in vivo* at two loci for generating knock-in rabbits (Song et al., 2016) but as with other compounds the effect appears to be cell-type specific and locus dependent (Xie et al., 2017; Zhang et al., 2017). Contradictory results for the putative DNA ligase IV inhibitor, Src7 (Riesenberg and Maricic, 2018; Song et al., 2016) have since been explained by incorrect chemical identity and lack of target specificity or potency (Greco et al., 2016). Compounds that can enhance the rate of HDR CRISPR efficiency *in vivo* are greatly needed (Pawelczak et al., 2017; Weidmann, 2018).

Competition between NHEJ and HR pathways exists (Her and Bunting, 2018). Two of the principal regulators are the epigenetic proteins 53BP1, which upregulates NHEJ; and BRCA1, which upregulates HR. The mechanisms of DSB repair pathway selection depends on the stage of the cell cycle, are complex and only partially understood (Chen et al., 2018).

Although NHEJ can occur at any stage of the cell cycle, BRCA1 negatively regulates 53BP1 and consequently NHEJ when the sister chromosomes are available during late S and G2 phases. 53BP1 is further negatively regulated during S and G2 phases by acetylation of its ubiquitylation-dependent recruitment (UDR) motif by CREB-binding protein controlled by HDAC2, thereby upregulating HR (Guo et al., 2017). Conversely, during G1 phase, when the sister chromosomes are not present, phosphorylated 53BP1 inhibits BRCA1 and HR, preventing DNA end resection of DSBs in a manner dependent upon its recruitment to histones and its downstream interaction with PTIP and RIF1 (Chen et al., 2018; Ward IM, 2003). NHEJ is a rapid mechanism of DSB repair, and can act within 30 minutes, whereas HR can take several hours (Her and Bunting, 2018; Panier and Boulton, 2014). Unsurprisingly, 53BP1 deficiency renders cells more susceptible to DNA damage including by ionising radiation (Panier and Boulton, 2014; Squatrito M, 2012). Somewhat unexpectedly, 53BP1 deficiency almost completely reverses the phenotype of BRCA1 deficiency, including in tumorigenesis, embryonic lethality and PARPi sensitivity (Bouwman et al., 2010; Bunting et al., 2010; Farmer et al., 2005; Panier and Boulton, 2014); effects that could be explained by partial restoration of HR.

53BP1 is a large multi-domain epigenetic reader protein that following RNF8 and RNF168 dependent ubiquitylation binds via its tandem tudor domain (TTD) to dimethylated lysine20 on the N-terminal tail of histone 4 (H4K20me2). The affinity of the interaction to a synthetic H4K20me2 peptide is ∼20 µM (Botuyan et al., 2006). It also binds via its UDR domain to a ubiquitin mark at Lysine 15 of histone H2A (Fradet-Turcotte A, 2013). Both histone modifications are key epigenetic marks for DSBs (Panier and Boulton, 2014).

53BP1 then recruits a wide set of DSB response proteins including RIF1, expand1 and PTIP (Bunting et al., 2010). Further recruitment of NHEJ effectors, including Ku70/80 complex, ultimately leads to DNA repair by DNA ligase IV (Panier and Boulton, 2014). Deleting 53BP1 *in vitro* leads to a ∼2-fold increase in HR (Orthwein et al., 2015), suggesting that 53BP1 might be a target for HDR CRISPR enhancement. By phage display screening, Canny et al. discovered a ubiquitin variant they named i53, which bound and inhibited the interaction of the TTD of 53BP1 with H4K20me2. Cells transfected with a plasmid expressing i53 showed 1.8-fold enhancement in HDR CRISPR efficiency at a single locus (Canny et al., 2018).

Similarly, Nambiar et al. showed that a RAD18 variant, e18 enhances HDR CRISPR *in vitro* approximately 2-fold by inhibiting 53BP1 localisation to DSBs (Nambiar et al., 2019). However, the use of such protein inhibitors expressed by plasmid transfection can be problematic *in vivo*, both from a plasmid delivery perspective, and because constitutive 53BP1 inhibition could interfere with normal development. Whereas chemical compounds can be transiently administered *in vivo* and do not require viral or nanoparticle delivery for gene therapy applications. We therefore tested a compound, EoHR (Figure 1A) that targets TTD of 53BP1 for 53BP1 inhibition and *in vivo* CRISPR applications.

**Figure 1:**
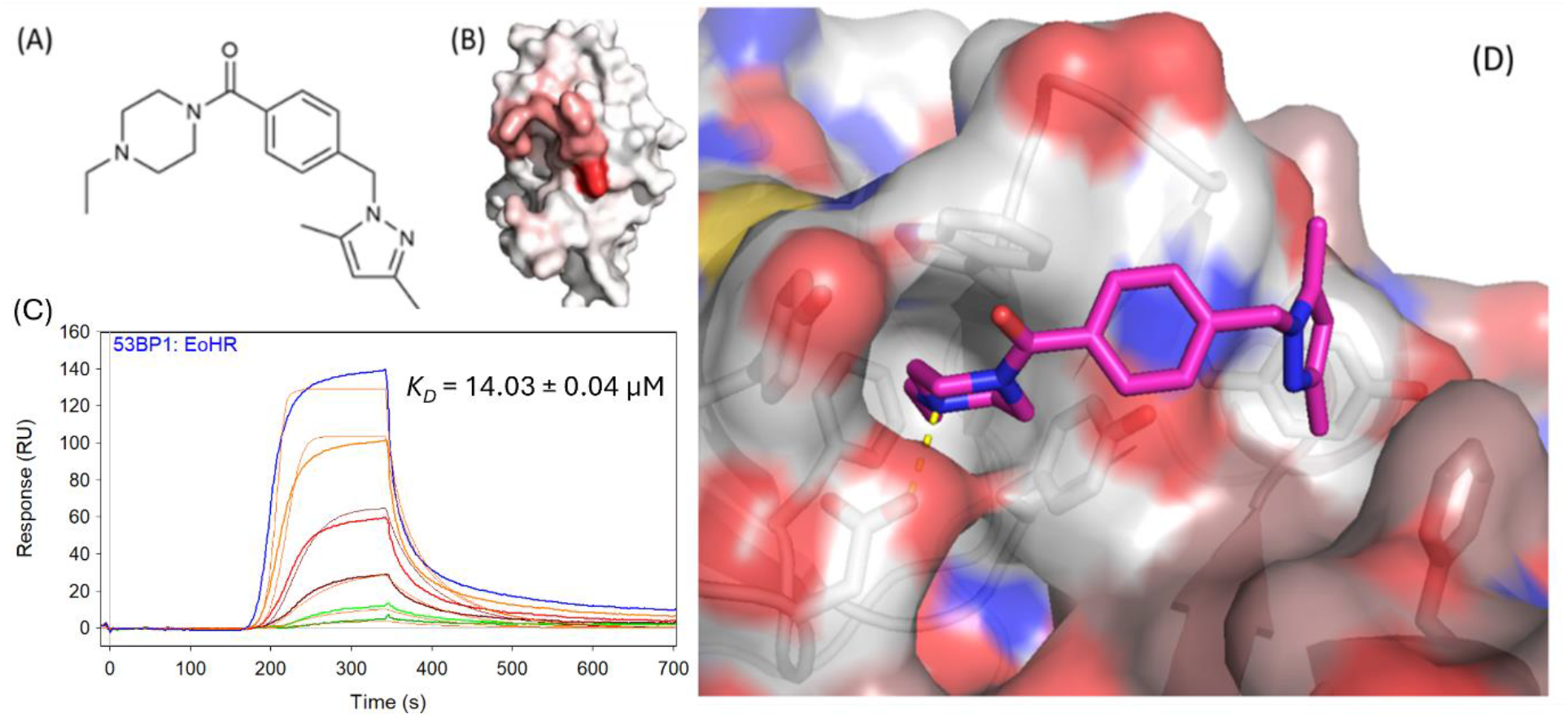
EoHR occludes the H4K20me2 binding site on 53BP1 TTD. **(A)** Molecular structure of EoHR. **(B)** Titration of EoHR into ^15^N-53BP1 TTD induces specific NMR chemical shift perturbations in the ^1^H,^15^N-HSQC spectrum of the protein that map to the H4K20me2 binding site. **(C)** SPR dose response sensorgrams of EoHR binding to immobilised 53BP1 TTD. **(D)** X-ray crystal structure of the complex reveals the ethyl piperazine warhead of EoHR occluding the H4K20me2 binding site on 53BP1 TTD. The hydrogen bond between D1521 and EoHR is shown in yellow dotted line. The crystal structure is rotated 90° anticlockwise with respect to the structure shown in (B).

### Results and Discussion EoHR binds 53BP1 TTD

EoHR binds the 53BP1 TTD with an affinity of 14 μM in our surface plasmon resonance (SPR) assay (Figure 1C). This is commensurate with the affinity of the 53BP1 TTD for the H4K20me2 peptide (Botuyan et al., 2006). Titration of EoHR into ^15^N-53BP1 TTD induces specific NMR chemical shift perturbations in the ^1^H,^15^N-HSQC spectrum of the protein that map to the H4K20me2 binding site (Figure 1B). The perturbation mapping is consistent with the binding pose observed in the 53BP1-EoHR X-ray crystal structure (Figure 1D). The ethyl piperazine of EoHR binds to the aromatic cage in one tudor domain where it makes a hydrogen bond with the key aspartate residue, D1521. The molecule bridges across to place the dimethyl pyrazole moiety of EoHR in the aromatic cage of the other 53BP1 tudor domain.

### EoHR inhibits 53BP1 localisation to double-strand DNA breaks

53BP1 contains two domains that are known to interact with histone marks for DSBs; the TTD that binds H4K20me2 (Botuyan et al., 2006), and the UDR domain which binds ubiquitinated lysine 15 on histone H2A (H2AK15ub) (Fradet-Turcotte A, 2013). Previous work demonstrated that mutation of either TTD H4K20me2 binding site residue D1521 or UDR residue L1619 is sufficient to abrogate 53BP1 accumulation at DNA repair foci (Fradet-Turcotte A, 2013; Zgheib et al., 2009). Here we tested whether EoHR can block 53BP1 accumulation at DSB repair sites using the eGFP-tagged 53BP1 construct (eGFP-53BP1) (Fradet-Turcotte A, 2013) in the DSB inducible via AsiSI (DIvA) cell system (Aymard F, 2017; Aymard F, 2014; Massip L, 2010). This cell system allows for the induction of approximately a hundred targeted DSBs by the AsiSI restriction enzyme, which translocates from the cytoplasm into the nucleus in response to its oestrogen receptor ligand-binding domain being activated by 4-hydroxytamoxifen (4OHT) addition. Prior to 4OHT treatment, nuclei exhibited diffuse eGFP-53BP1 fluorescence, with a few intensely fluorescent DNA repair foci. In the absence of EoHR, induction of the DNA endonuclease caused 53BP1 to accumulate at DNA DSBs over a period of 0.5 to 2 hours, with typically 15-25 foci forming per cell. In the presence of EoHR the formation of 53BP1-GFP foci was inhibited in a dose-dependent manner (Figure 2). The IC_50_ estimated at 2 hours post OHT addition was 13 ± 5 µM. This demonstrates that EoHR can recapitulate the phenotype of the D1521 mutant (Fradet-Turcotte A, 2013; Zgheib et al., 2009) and block 53BP1 accumulation at DSB repair sites.

**Figure 2:**
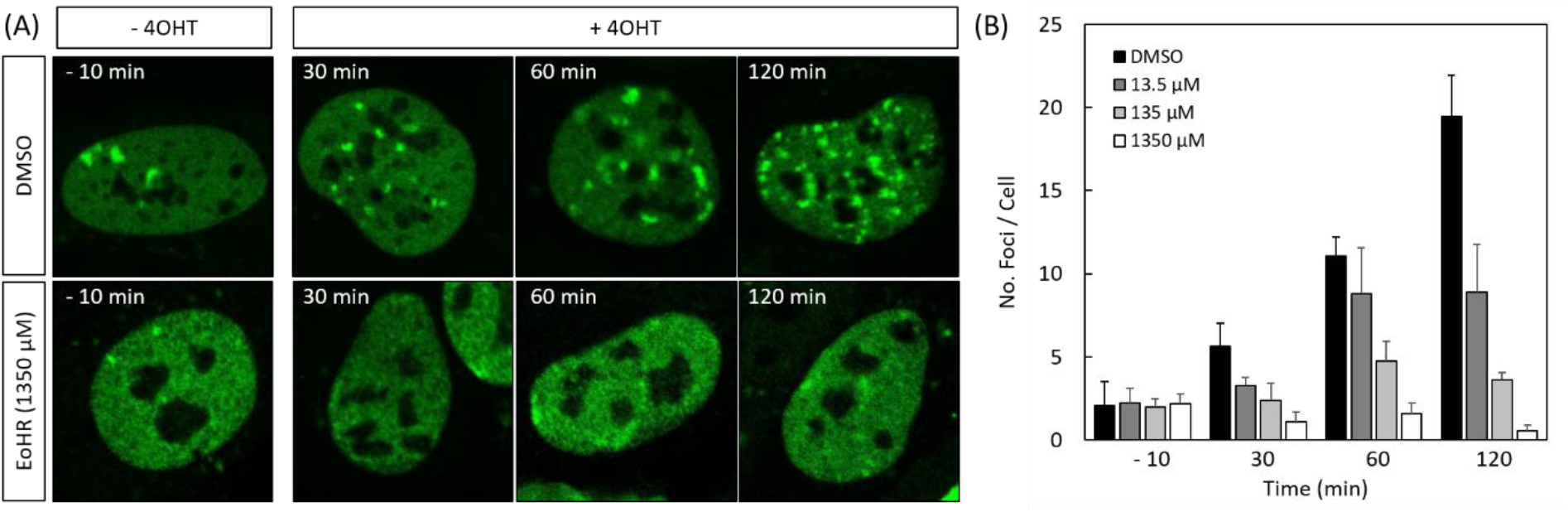
The effect of EoHR on recruitment of 53BP1 to DSBs. **(A)** eGFP-53BP1 fluorescence in the nuclei of cells before and after induction of DNA double-strand breaks via 4OHT treatment in the presence and absence of EoHR (1350 µM). **(B)** Dose response titration of EoHR. The histogram depicts the mean number of eGFP-53BP1 foci per nucleus before and after DSB induction. Errors bars indicate standard error of the mean.

No evidence of cell toxicity was observed in these DlvA cells at any concentration up to 1350 µM. HCT116 cells cultured for 3 days in the presence of EoHR up to 1000 μM showed only mild growth retardation at the highest dose (SI Figure 2). These results support the use of EoHR as a tool compound for investigating 53BP1 inhibition at DSBs.

### EoHR enhances HDR CRISPR efficiency for the creation of transgenic mice

Mouse and human 53BP1 TTDs share 100% sequence identity. Thus, we proposed that at inhibitory concentrations, EoHR should outcompete 53BP1 binding to H4K20me2 modified histones at DSBs induced by Cas9, inhibiting NHEJ and favouring HDR in CRISPR experiments.

Preliminary experiments with naked DNA showed that EoHR at concentrations up to 5 mM did not interfere with Cas9 nuclease activity (see SI, Figure S2). To test the efficacy of EoHR in enhancing HDR CRISPR we trialled the compound in 17 independent and unique HDR projects, all of which involved the generation of insertions using HDR CRISPR. Insert sizes ranged from 652 to 2111 base pairs (Table 1) and each insertion involved a distinct locus (17 loci in total on 11 chromosomes). The HDR CRISPR protocol involved the microinjection of sgRNA, Cas9 and the HDR DNA repair template into a pronucleus of a fertilised mouse oocytes at the single-cell stage (Harms et al., 2014) with or without 5 mM EoHR added to the microinjection buffer. Assuming full cellular diffusion and an oocyte volume of ∼180 picolitres (Sorensen and Wassarman, 1976), injection of 4-8 picolitres will result in a final *in vivo* concentration of 110-220 µM, which is ∼10 times its IC_50_ and should provide ∼95% inhibition. Microinjected eggs were then transferred to the oviducts of pseudopregnant females.

**Table 1:**
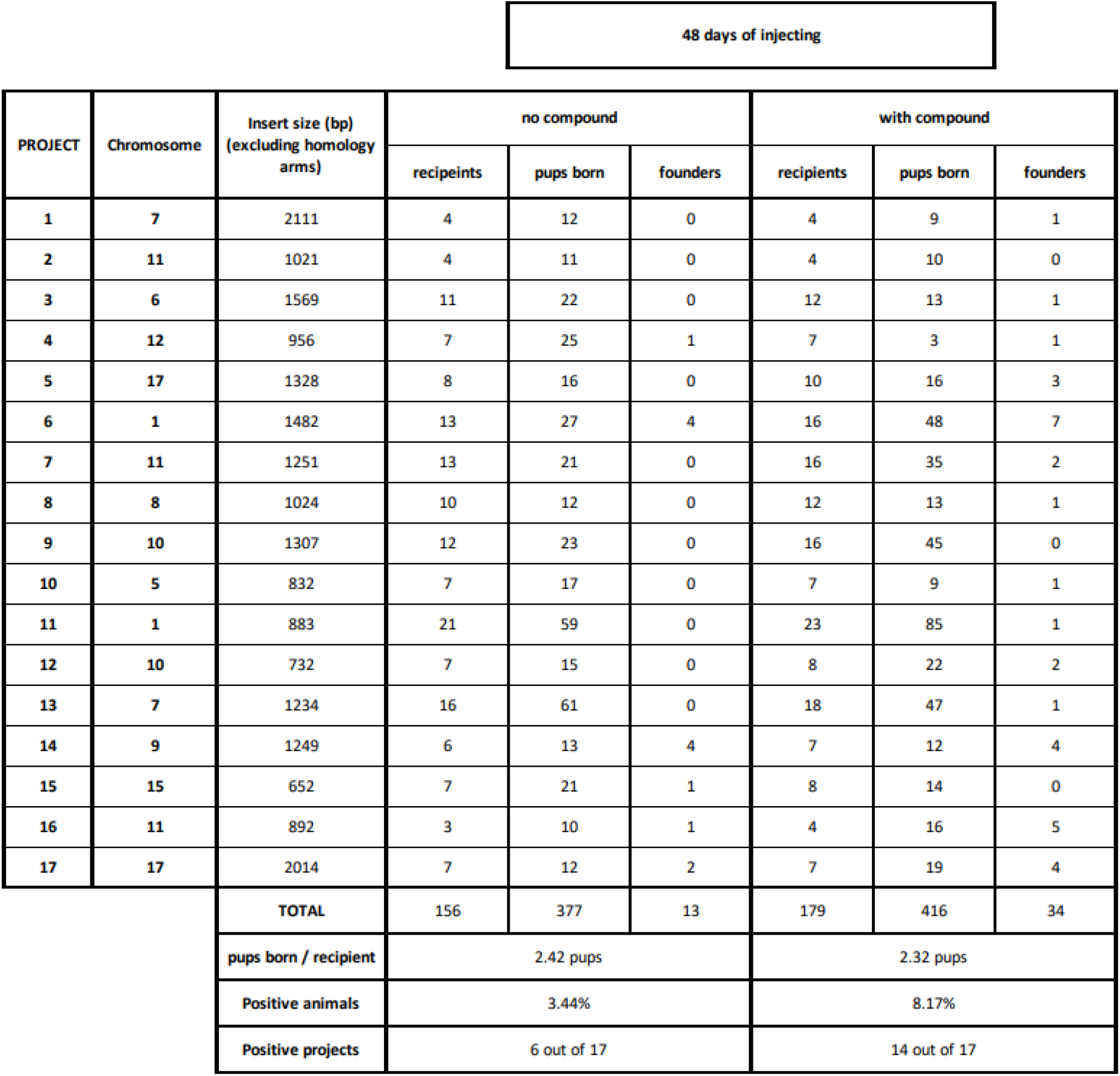
Summary data for the HDR CRISPR projects investigated. The 17 projects required DNA repair templates that were unique and ranged in size from 652 to 2111 base pairs, with each HDR CRISPR insertion targeting a distinct locus. The chromosome targeted is given. In total 48 experiments were conducted with and without the addition of EoHR. The table depicts the number of recipient females for zygote implantation, live pups per litter and founder pups per project. The principal statistics are given; average number of live pups per litter, the founder success rate, proportion of positive projects.

| 48 days of injecting |  |  |  |  |  |  |  |  |
| --- | --- | --- | --- | --- | --- | --- | --- | --- |
| PROJECT | Chromosome | Insert size (bp)<br>(excluding homology arms) | no compound |  |  | with compound |  |  |
|  |  |  | recipients | pups born | founders | recipients | pups born | founders |
| 1 | 7 | 2111 | 4 | 12 | 0 | 4 | 9 | 1 |
| 2 | 11 | 1021 | 4 | 11 | 0 | 4 | 10 | 0 |
| 3 | 6 | 1569 | 11 | 22 | 0 | 12 | 13 | 1 |
| 4 | 12 | 956 | 7 | 25 | 1 | 7 | 3 | 1 |
| 5 | 17 | 1328 | 8 | 16 | 0 | 10 | 16 | 3 |
| 6 | 1 | 1482 | 13 | 27 | 4 | 16 | 48 | 7 |
| 7 | 11 | 1251 | 13 | 21 | 0 | 16 | 35 | 2 |
| 8 | 8 | 1024 | 10 | 12 | 0 | 12 | 13 | 1 |
| 9 | 10 | 1307 | 12 | 23 | 0 | 16 | 45 | 0 |
| 10 | 5 | 832 | 7 | 17 | 0 | 7 | 9 | 1 |
| 11 | 1 | 883 | 21 | 59 | 0 | 23 | 85 | 1 |
| 12 | 10 | 732 | 7 | 15 | 0 | 8 | 22 | 2 |
| 13 | 7 | 1234 | 16 | 61 | 0 | 18 | 47 | 1 |
| 14 | 9 | 1249 | 6 | 13 | 4 | 7 | 12 | 4 |
| 15 | 15 | 652 | 7 | 21 | 1 | 8 | 14 | 0 |
| 16 | 11 | 892 | 3 | 10 | 1 | 4 | 16 | 5 |
| 17 | 17 | 2014 | 7 | 12 | 2 | 7 | 19 | 4 |
| TOTAL |  |  | 156 | 377 | 13 | 179 | 416 | 34 |
| pups born / recipient |  |  | 2.42 pups |  |  | 2.32 pups |  |  |
| Positive animals |  |  | 3.44% |  |  | 8.17% |  |  |
| Positive projects |  |  | 6 out of 17 |  |  | 14 out of 17 |  |  |

In total, across the 17 projects, 793 pups were born and genotyped (Table 1). Injection of the compound showed no evidence of toxicity to the developing foetuses as assessed by the number of live births per litter (EoHR 2.32 ± 0.95 vs control 2.42 ± 0.73, n = 335 litters, p = 0.42, Figure 3C).

**Figure 3:**
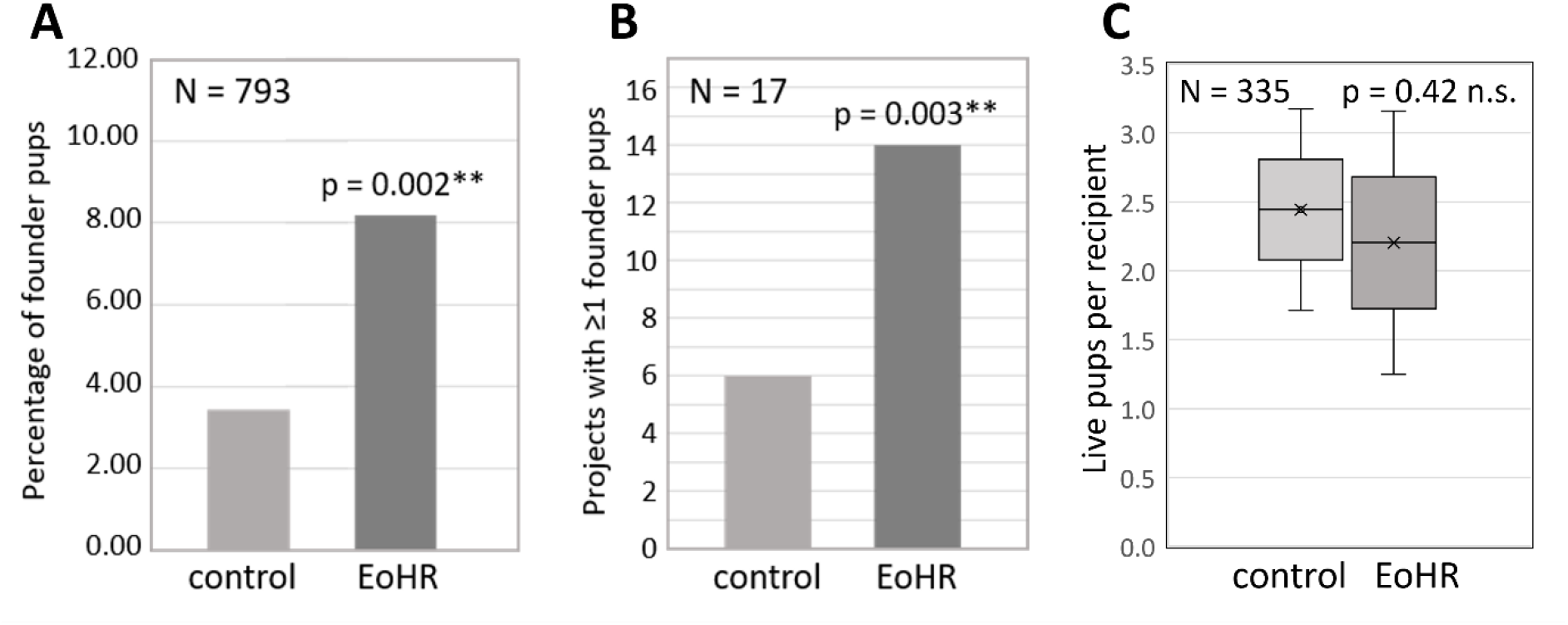
Effects of EoHR on knock-in rates *in vivo*. **A** Efficiency of Cas9-mediated knock-in mouse pups genotyped at birth (N = 793 pups). **B** The number of projects having at least one knock-in pup (N = 17 projects). **C** The number of live pups born per mouse dam.

Genotyping revealed that EoHR addition to the standard microinjection buffer resulted in a 2.4-fold enhancement of HDR CRISPR efficiency combined across all projects (EoHR 8.17% vs control 3.44%, N=793 pups genotyped, p=0.002**) (Figure 3A, Table 1). Notably, an increase in HDR gene targeting efficiency of ∼2-fold was the effect size observed in 53BP1-null cells (Orthwein et al., 2015) and for 53BP1 inhibition by i53 and e18 *in vitro* (Canny et al., 2018; Nambiar et al., 2019). The 2.4-fold effect in this study is therefore likely to approximate the maximal effect size achievable through 53BP1 inhibition and is consistent with the near-complete inhibition of 53BP1-GFP accumulation at DNA repair foci that we observed in the presence of saturating concentrations of EoHR.

The success of an HDR CRISPR project ultimately only requires the generation of a single viable knock-in founder pup. Hence each project was concluded once a viable founder pup was established. Conceptually, if a compound were to only enhance the success rate of more straightforward HDR CRISPR projects it would be of limited utility. We therefore assessed the project success rate in the presence and absence of EoHR injection. Project success rate was similarly 2.3-fold enhanced by EoHR (14 vs 6 projects, n = 17, p=0.003*) (Figure 3B, Table 1) mirroring the cumulative HDR CRISPR efficiency enhancement. The implication is that EoHR enhances the likelihood of successful HDR CRISPR in mice irrespective of the locus that is targeted.

Acquiring a transgenic mouse model using CRISPR knock-in typically takes 6-12 months; largely because the high HDR failure rates typically require multiple experimental cycles. Breeding and housing the mice is the most expensive and rate limiting step.

Consequently, adding EoHR to the standard protocol will double the efficiency with which knock-in mice can be generated. This will substantially increase the throughput of transgenic mouse providers and should reduce the time and cost for scientists to obtain a knock-in mouse.

### EoHR is well tolerated in neonatal mice but did not increase gene insertion in liver

Many metabolic diseases are caused by mutations or deletions in enzymes produced in the liver. These are a relatively attractive targets for developing knock-in CRISPR therapies as partial restoration of circulating enzyme levels can ameliorate disease (Adlat et al., 2023). We therefore tested the ability of EoHR to enhance gene insertion efficiency in neonatal mice using liver-specific adeno-associated viral (AAV) vectors to create an albumin-GFP fusion protein under regulation of the albumin promoter (De Caneva et al, 2019). Intraperitoneal injections of EoHR were given daily on days 1, 2 and 3 postpartum at 75 mg/kg. At day 19 when livers when animals were euthanised and livers harvested there was no discernible difference in the size or condition of the mice or their livers between the EoHR and control groups. However, GFP gene incorporation into hepatocytes in the EoHR group (3.6% ± 2.2%) was not significantly increased compared to controls (2.5% ± 1.3%) as assessed by student’s t-test (p = 0.10 n.s.).

## Conclusions

EoHR provides researchers with a low-toxicity compound for exploring the various roles of 53BP1 in DNA repair pathways. We have shown that EoHR enhanced HDR CRIPR for creating transgenic mice. As the 53BP1 amino acid sequence of 53BP1 TTD is also 100% conserved in all vertebrates for which gene sequencing data is available, including in agriculturally important animals such as pigs and cows, similar increases in gene insertion efficiency can reasonably be expected with EoHR. Other biotech applications may also benefit from the use of EoHR. For instance, reprogramming cells to pluripotency has recently been shown to improve in the absence of 53BP1 (Georgieva et al. 2024).

It seems likely that higher affinity 53BP1 inhibitors with good pharmacokinetic and toxicology profiles will need to be developed for *in vivo* HDR gene therapy applications. In this regard, the low toxicity and relatively small size of EoHR makes it an attractive lead for drug development.

## Methods

### 53BP1 expression and purification

The pET15 53BP1 TTD expression construct was transformed into Rosetta BL21(DE3)pLysS competent cells (Novagen). Cells were grown at 37°C with shaking until the OD600 reached ∼0.6-0.8 at which time the cells were harvested by centrifugation and resuspended in minimal media containing ^15^NH_4_Cl 1g/L and glucose 4g/L. After growth was re-established, the temperature was lowered to 18°C and expression induced by adding 0.5mM IPTG and continuing shaking overnight. Cells were harvested by centrifugation.

His-tagged 53BP1 TTD was purified by resuspending thawed cell pellets in 30ml of lysis buffer (50mM sodium phosphate pH 7.2, 50mM NaCl, 30mM imidazole, 1 x EDTA free protease inhibitor cocktail (Roche) per litre of culture). Cells were lysed on ice by sonication. The cell lysate was clarified by centrifugation and purified by FPLC using Ni-NTA (HisTrap FF column, GE Healthcare) followed by size exclusion chromatography (HiLoad 26/60 Superdex 75, GE Healthcare) that had been preequilibrated with 1.2 column volumes of NMR buffer (25mM sodium phosphate buffer pH 7.2, 100mM NaCl).

### Surface Plasmon Resonance

SPR experiments were performed on a SensiQ Pioneer FE (Sartorius) at 25°C, with running buffer 10 mM sodium phosphate pH 7.4, 150 mM NaCl, 3 mM KCl, 50 μM EDTA, 5% DMSO, 0.005% Tween-20. His_6_-tagged 53BP1 TTD was captured on an NHS/EDC activated Sartorius HisCap chip in the presence of 20 µM EoHR. EoHR affinity was determined using OneStep gradient injections of EoHR in 3-fold maximum concentrations from 400 nM to 100 μM. Sensorgrams were fitted using the kinetic model with a mass transport parameter in QDat software (Sartorius).

### NMR spectroscopy

The NMR sample comprised 70 μM ^15^N-53BP1 TTD in NMR buffer (25 mM sodium phosphate pH 7.4, 50 mM NaCl, 10% D2O). Spectra were acquired at 25°C on a Bruker Avance III 600 MHz spectrometer equipped with a TXI-cryoprobe and processed with Topspin 3.5. ^1^H,^15^N-HSQC and ^1^H 1D spectra were acquired for ^15^N-53BP1 TTD in its apo state and in the presence of EoHR 360 µM and the chemical shift perturbations were analysed using SPARKY with reference to the published assignments (Charier et al., 2004).

### Crystallisation

Purified 53BP1 (10 mg/mL) in 20 mM Tris pH7.5, 200 mM NaCl was crystallized by mixing 1 µL of protein solution containing EoHR (11.4 mM) with 1 µL of reservoir solution containing 20% PEG3350 and 0.2M Mg(NO_3_)_2_ and the best crystals were obtained by vapor diffusion technique at 20°C in sitting drops. For cryoprotection, the crystals were soaked briefly in the reservoir solution supplemented with 15% ethylene glycol (v/v) before flash freezing in liquid N_2_.

### X-ray Data Collection and Structure Determination

X-ray diffraction data for 53BP1+EoHR was collected at 100 K at beam line 08ID-1 of CMCF at Canadian Light Source. The data set was processed using the HKL-30001 suite (Otwinowski and Minor, 1997). The structures of 53BP1 + EoHR was solved by molecular replacement using PHASER2 (McCoy et al., 2007) with PDB entry 4RG2 as search template. Geometry restraints for the compound refinement were prepared with GRADE3 (O. S. Smart, 2011) developed at Global Phasing Ltd. Graphics program COOT4 (P. Emsley 2010) was used for model building and visualization. Restrained refinement and validation using REFMAC5 (Murshudov et al., 1997), and MOLPROBITY6 (Chen VB, 2010), respectively. The structure was deposited in the Protein Data Bank 6VIP (The et al., 2020).

### Cell Culture and Treatment

DIvA cells (originally provided by Gaëlle Legube, LBCMCP, CNRS, Toulouse, France) were grown in DMEM (Lonza) supplemented with 10% bovine growth serum (Gibco), 1 x Pen-Strep (Lonza) and 1 µg/ml puromycin (ThermoFisher Scientific) at 37°C in 5% CO _2_. The DIvA cells were plated 24 h before experiments onto 8 well ibidi glass bottom dishes and transiently transfected with eGFP-53BP1 (Fradet-Turcotte A, 2013) (Addgene #60813) via use of Lipofectamine 3000 according to the manufacturer’s protocol. Transfected DIvA cells are pre-incubated with EoHR (13.5, 135 and 1350 μM) for 2 hours. DSB induction was then performed in the absence and presence of EoHR via treatment of DIvA cells with 4OHT (300 nM) (Aymard F, 2017; Aymard F, 2014; Massip L, 2010). Cells were then fixed by 4% PFA for 15 minutes at room temperature and rinsed with PBS 3 times before imaging.

### Fluorescence Microscopy and Analysis

All microscopy measurements were performed on an Olympus FV3000 laser scanning microscope. A 60x/1.2NA water immersion objective was used for all experiments and the cells were imaged at 37°C in 5% CO_2_. eGFP-53BP1 was excited by a solid-state laser diode operating at 488 nm and the fluorescence signal of eGFP was directed through a 488/561 dichroic mirror to a H7422P of Hamamatsu photomultiplier detector fitted with a GFP 520/25 nm bandwidth filter. The number of eGFP-53BP1 foci per DiVA nucleus imaged was quantified in CellProfiler.

### Cas9 inhibition test

Recombinant *S. pyogenes* Cas9 nuclease, 0.5ug, (IDT) and sgRNA, 20 ng, in Cas9 nuclease reaction buffer (NEB) were incubated for 10 minutes at 25°C in the absence or presence of 1 mM EoHR (1% DMSO) or 5 mM EoHR (5% DMSO). The dsDNA template, 100 ng, was then added to the samples and incubated for 60 minutes at 37°C with water and 5% DMSO controls. Samples were analysed on 2% agarose gel with ethidium bromide under UV light.

### EoHR synthesis

#### EoHR for cell culture and SPR was synthesised as follows

*4-((3,5-dimethyl-1H-pyrazol-1-yl)methyl)benzoic acid*: 3,5-dimethylpyrazole (500 mg, 5.2 mmol) was dissolved in DMF (15 mL) and sodium tert-butoxide (1 g, 10.4 mmol) was added to this solution, and the mixture was stirred at room temperature for 15 minutes. A solution of 4-(bromomethyl)benzoic acid (214 mg, 1 mmol), in DMF (20 mL) was then added dropwise to the reaction mixture and stirred for 30 min. The solvent was then removed under vacuum, and the residue dissolved in H_2_O (50 mL). The solution was acidified to pH 5 with aqueous HCl (2 M), and the resulting precipitate was filtered off to afford the title compound (0.65 mg, 82%) as a white powder. This product was subjected to the next step without further purification. ^1^H NMR (300 MHz, CDCl_3_) δ 12.61 (s, 1H, COOH), 8.00 (d, *J* = 8.2 Hz, 2H, Ar-H), 7.13 (d, *J* = 8.2 Hz, 2H, Ar-H), 5.89 (s, 1H, Het-H), 5.35 (s, 2H, Ar-CH_2_), 2.28 (s, 3H, Het-CH_3_), 2.14 (s, 3H, Het-CH_3_). ^13^C NMR (75 MHz, CDCl_3_) δ 170.4, 148.0, 142.6, 140.0, 130.7, 129.6, 126.7, 106.1, 52.1, 13.3, 11.2.

*(4-((3,5-dimethyl-1H-pyrazol-1-yl)methyl)phenyl)(4-ethylpiperazin-1-yl)methanone*: The benzoic acid (0.47 g, 2.05 mmol) dissolved in 60 mL of DMF, then HATU (0.90 g, 2.36 mmol, 1.15 equiv.) and DIPEA (0.53 g, 4.10 mmol, 2.00 equiv.) were added. The reaction mixture was stirred at room temperature for 30 minutes under nitrogen, then *N*-ethylpiperazine (0.47 mg, 4.10 mmol, 2.00 equiv.) was added. This mixture was stirred overnight, then solvent was removed under reduced pressure. The mixture was redissolved in 100 mL of water and saturated sodium bicarbonate to pH 8. This mixture was extracted with DCM (4 x 50 mL), which was separated then washed with brine (3 x 50 mL). The DCM extract was dried with sodium sulfate, then the solvent was removed under reduced pressure to give a yellow oil. This was separated using column chromatography (4% methanol in DCM with an additional 1% ammonium hydroxide, dried over activated silica) to give the product as yellow oil (0.56 mg, 84%). ^1^H NMR (300 MHz, CDCl_3_) δ 7.34 (d, *J* = 8.0 Hz, 2H, Ar-H), 7.07 (d, *J* = 8.0 Hz, 2H, Ar-H), 5.86 (s, 1H, Het-H), 5.23 (s, 2H, Ar-CH_2_), 3.85-3.51 (m, 4H, CH_2 &_ NCH_2_), 2.55-2.53 (m, 6H, NCH_2_), 2.24 (s, 3H, Het-CH_3_), 2.15 (s, 3H, Het-CH_3_), 1.15 (s, 3H, NCH_2_CH_3_). Δ ^13^C NMR 170.1, 148.2, 139.7, 139.5, 134.5, 127.8, 126.8, 52.4, 52.2, 13.7, 11.2. HRMS (EI) calculated for C19H26N4O *m/z* 326.211 [M]^+^; observed: 326.2.

EoHR for mouse studies was purchased from Enamine Pty Ltd, catalogue no. EN300-719578, CAS 1281035-76-4. Identity and purity were confirmed in-house by NMR spectroscopy.

### Transgenic Mouse HDR CRISPR

The protocol used was the standard protocol (Harms et al., 2014). Briefly, upon microinjection days the number of fertilised mouse eggs was equally divided. The control injection solutions contained containing recombinant *S. pyogenes* Cas9 nuclease, 10 – 30 ng/µl (IDT), the crRNA/tracrRNA complex, 10 – 30 ng/µl (IDT), and the ssDNA repair template for the project (10 – 30 ng/µL) in microinjection buffer (10 mM Tris-HCl pH 7.4, 1mM EDTA, 100mM NaCl). The experimental injection solutions were the same except with the addition of EoHR 5mM in 5% DMSO. Sequencing of PCR mouse tail DNA reactions verified the correct integration of the genetic insert using forward and reverse primers external to the homology arms and uniquely designed for each insert. Toxicity was assessed by the number of live births per litter using the Student’s t-test. Statistical tests for HDR CRISPR efficiency and project success rate were derived using the normal approximation for the binomial distributions of the observed frequencies (Zaiontz, 2013). For the full statistical derivation see supplementary information.

### HDR CRISPR in live neonatal mice

All mouse procedures described here were conducted in accordance with the recommendations of the National Health and Medical Research Council Australia Code of Practice for the Care of Experimental Animals and protocols approved by the RMIT University Animal Ethics Committee. Pregnant Swiss-Webster mice were obtained from the Animal Resources Centre, WA, Australia. On arrival at the RMIT University Animal Facility, they were housed at 22°C, 40-60% humidity, 12 h light/dark cycle (7 am to 7 pm) and provided with ad libitum pelleted standard rat chow and water.

On day 1 or 2 after birth, the dams were temporarily removed from the home cage and 59 pups from 8 litters were weighed and then individually chilled on crushed wet ice for ∼1 minute to partially anaesthetise them. The two AAV8 vectors were mixed to give a final concentration of 8.0 vg/mouse (rAAV8-donor-eGFP) and 4.0 vg/mouse (rAAV8-SaCas9-sgRNA8). The mixture was then injected into the cranial vein behind the ear bud using a 30G insulin needle at an injection volume of 15 µL. On the same day, 29 of the pups were given the EoHR compound i.p. (75mg/kg/day in 5% DMSO; 20 µL injection volume), with the remainder receiving vehicle control. Following injection, pups were warmed until moving as normal and then returned to the litter in the home cage. When the entire litter was injected, the dam was returned to the cage and observed until suckling normally. The EoHR compound was administered again at the same dose on days on the next two consecutive days.

Mice were killed on day 19 after injection by cervical dislocation, livers extracted and immersion-fixed in 4% paraformaldehyde overnight before being stored in 20% sucrose in phosphate-buffered saline. Liver samples were cut into 30 µm sections on a cryostat, sections were slide-mounted, stained with DAPI and coverslips applied. Images were acquired on Olympus BX60 fluorescence microscope and analysed using ImageJ (NIH) to determine the percentage of cells expressing GFP. Samples were analysed blinded with at least three sections per liver being quantitated.

## Acknowledgements

The authors would like to thank Dr. Jieqiong Lou and Dr Elizabeth Hinde (University of Melbourne) for performing the experiments with the DSB inducible via AsiSI cell system (DIvA), and Gaëlle Legube and Dr. Thomas Clouaire for providing the cells. Dr Lorena Zentilin and Andrés Muro kindly provided the AAV vectors for the juvenile mice experiments. The transgenic mouse experiments were performed at Monash Genome Modification Platform. The 53BP1 plasmid was provided by Dr Ingerman UNC and Dr Brown SGC Toronto solved the 53BP1/EoHR crystal structure.

## Author contributions

SS, MM, AS, juvenile mouse experiments; SM, FOD, synthesised EoHR and performed SPR experiments; AL, BH, FR, injection of oocytes; AR, AG, AJD cell toxicity experiments. SH, expressed protein, performed SPR and NMR experiments, designed and managed project, coordinated manuscript.

## Notes

### Competing Interest Statement

The authors have declared no competing interest.

