## Supplementary Information for "Improved gene targeting in vivo using EoHR, a small molecule inhibitor of 53BP1"

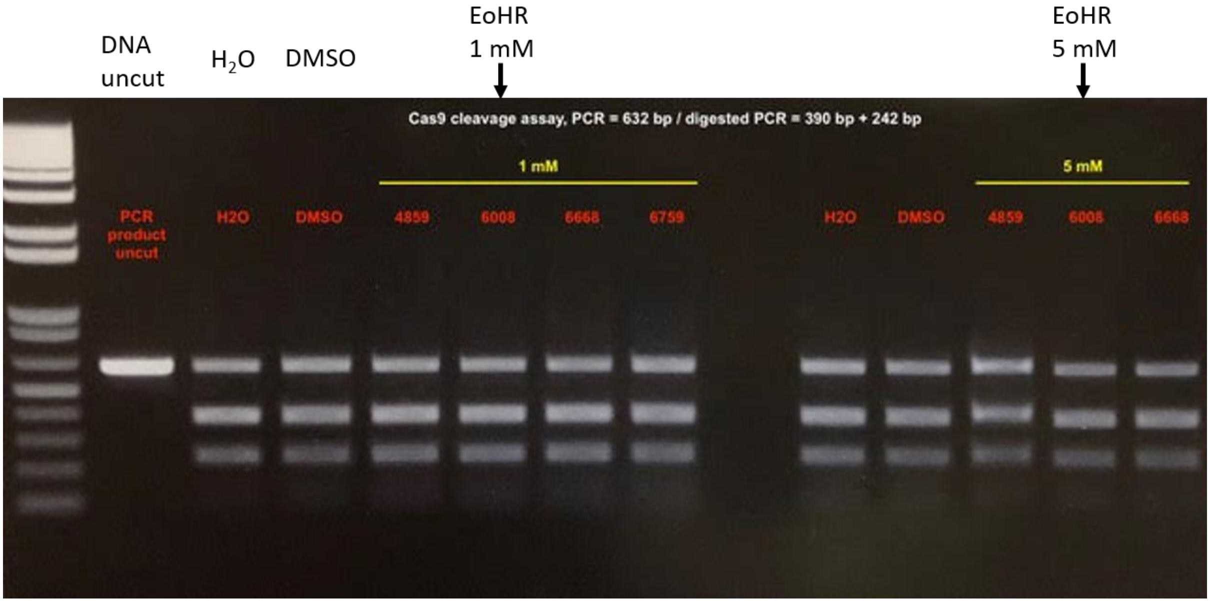


**Supplementary Figure 1. Cas9 DNA endonuclease activity in the presence of EoHR at 1 and 5 mM.** Recombinant *S.pyogenes* Cas9 nuclease and sgRNA were incubated for 10 minutes at 25 ˚C in the absence or presence of 53BP1-binding compounds including 1 mM EoHR (1% DMSO) or 5 mM EoHR (5% DMSO). A dsDNA template was then added to the samples and incubated for 60 minutes at 37 ˚C with water and 5% DMSO controls. Samples were analysed on 2% agarose gel, with ethidium bromide under UV light.

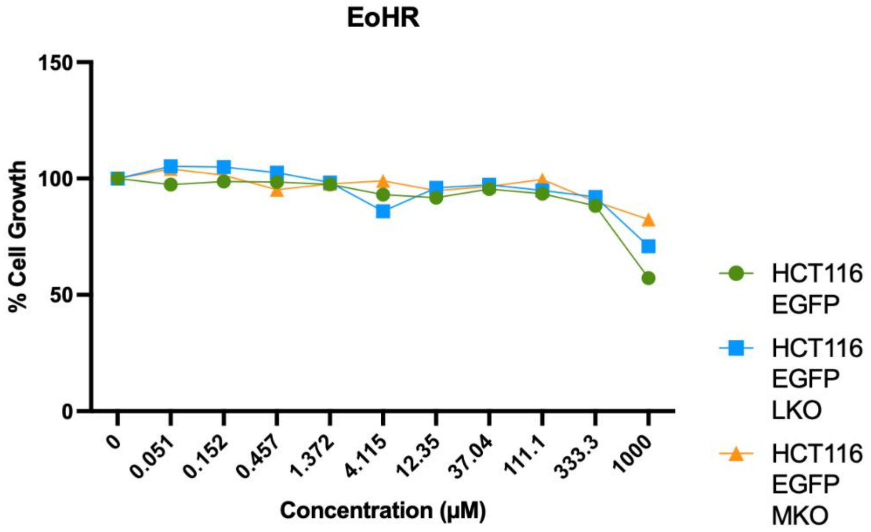


**Supplementary Figure 2. HCT cell growth in the presence of EoHR.** HCT116 cells were cultured for 3 days in the presence of EoHR up to 1000 μM.

**Statistical analysis of knock-in mouse experiment**

Adapted with permission from; Zaiontz C. (2019) Real Statistics Using Excel. www.real-statistics.com

Theorem: Let $x_1$ and $x_2$ be random variables with proportional distributions with mean $\pi_1$ and $\pi_2$ respectively. Let $p_1$ be the proportion of successes in $n_1$ trials of the first distribution and let $p_2$ be the proportion of successes in $n_2$ trials of the second distribution. When the number of trials $n_1$ and $n_2$ are sufficiently large, usually when $n_i \pi_i \geq 5$ and $n_i(1 - \pi_i) \geq 5$, the difference between the sample proportions $p_1 - p_2$ will be approximately normal with mean $\pi_1 - \pi_2$ and standard deviation

$$\sqrt{\frac{\pi_1(1-\pi_1)}{n_1} + \frac{\pi_2(1-\pi_2)}{n_2}}$$

Normality criteria

For total founder pups the binomial distribution closely approximates a normal distribution (ie. criteria >>5) for founder pups with sample sizes control = 377 and experiment = 416 and p(control) = 0.034 and p(experiment) = 0.082. For projects the normality criteria are >5 for n = 17 and p(control) = 0.35 and p(experiment) = 0.82. Therefore, the following proof and statistical test can be applied.

Proof: Based on Theorem 2 of the binomial distribution, $x_i$ has approximately the distribution

$$N(\pi_i, \sqrt{\pi_i(1-\pi_i)/n_i})$$

Since $x_1$ and $x_2$ are independently distributed, by the linear transformation property of the normal distribution, $x_1 - x_2$ has distribution

$$N(\pi_1 - \pi_2, \sqrt{\pi_1(1-\pi_1)/n_1 + \pi_2(1-\pi_2)/n_2})$$

Statistical test

We now test the following null hypothesis: $H_0$: $\pi_1 = \pi_2$. Assuming the null hypothesis is true, by Theorem 2, $x_1 - x_2$ will be approximately normal with mean $\pi_1 - \pi_2 = 0$ and standard deviation

$$\sqrt{\frac{\pi_1(1-\pi_1)}{n_1} + \frac{\pi_2(1-\pi_2)}{n_2}}$$

We use a two-tail test with a significance cut-off at >0.975 using the normal distribution function in Microsoft Excel.

HDR CRISPR efficiency, n = 793 pups

Inverse p-value = NORMDIST(0.04725, 0, 0.01664, TRUE) = 0.998

p-value = 0.002**

Project success rate, n = 17 projects

Inverse p-value = NORMDIST(0.471, 0, 0.174, TRUE) = 0.997

p-value = 0.003**

Thus, in both cases we reject the null hypothesis, and conclude there is a significant difference between experiment and control for HDR CRISPR efficiency rate and project success rate.
